# Environmental perturbations lead to extensive directional shifts in RNA processing

**DOI:** 10.1101/119974

**Authors:** A. L. Richards, D. Watza, A. Findley, A. Alazizi, X. Wen, A. A. Pai, R. Pique-Regi, F. Luca

## Abstract

Environmental perturbations have large effects on both organismal and cellular traits, including gene expression, but the extent to which the environment affects RNA processing remains largely uncharacterized. Recent studies have identified a large number of genetic variants associated with variation in RNA processing that also have an important role in complex traits; yet we do not know in which contexts the different underlying isoforms are used. Here, we comprehensively characterized changes in RNA processing events across 89 environments in five human cell types and identified 15,300 event shifts (FDR = 15%) comprised of eight event types in over 4,000 genes. Many of these changes occur consistently in the same direction across conditions, indicative of global regulation by trans factors. Accordingly, we demonstrate that environmental modulation of splicing factor binding predicts shifts in intron retention, and that binding of transcription factors predicts shifts in AFE usage in response to specific treatments. We validated the mechanism hypothesized for AFE in two independent datasets. Using ATAC-seq, we found altered binding of 64 factors in response to selenium at sites of AFE shift, including ELF2 and other factors in the ETS family. We also performed AFE QTL mapping in 373 individuals and found an enrichment for SNPs predicted to disrupt binding of the ELF2 factor. Together, these results demonstrate that RNA processing is dramatically changed in response to environmental perturbations through specific mechanisms regulated by trans factors.

**Author Summary:** Changes in a cell’s environment and genetic variation have been shown to impact gene expression. Here, we demonstrate that environmental perturbations also lead to extensive changes in alternative RNA processing across a large number of cellular environments that we investigated. These changes often occur in a non-random manner. For example, many treatments lead to increased intron retention and usage of the downstream first exon. We also show that the changes to first exon usage are likely dependent on changes in transcription factor binding. We provide support for this hypothesis by considering how first exon usage is affected by disruption of binding due to treatment with selenium. We further validate the role of a specific factor by considering the effect of genetic variation in its binding sites on first exon usage. These results help to shed light on the vast number of changes that occur in response to environmental stimuli and will likely aid in understanding the impact of compounds to which we are daily exposed.

## Introduction

Variation in gene expression has long been associated with cellular and organismal phenotypes. For example, studies have found that gene expression in blood and bronchial epithelial cells differs among individuals with asthma [1, 2, 3, 4]. Such differences in gene expression occur in specific cellular pathways, such as the glucocorticoid response pathway [1, 5, 6, 7], leading to the general usage of glucocorticoids to treat asthma. These studies, and others, have demonstrated that variation in gene expression plays a role in complex traits and cellular responses [8, 9, 10, 11, 12]. More recently, however, researchers have begun to assess the impact of alternative mRNA isoform usage on phenotypes. Previous studies have found that RNA processing, leading to differential isoform usage, is different in certain diseases such as Alzheimer’s disease and several forms of cancer [13, 14, 15, 16, 17]. Furthermore, studies have identified global shifts in exon usage associated with developmental or diseased cellular states. For instance, shorter 3’ untranslated region (UTR) isoforms are prevalent in proliferating or cancerous cells [18, 19]. Cancer is also associated with increased retention of introns [20, 21].

Li *et al.* recently identified genetic variants associated with inter-individual variation in mRNA splicing and identified almost 2,900 splicing Quantitative Trait Loci (QTLs). Further, they showed that splicing QTLs are also enriched for genetic variants associated with several complex traits in Genome-Wide Association Studies (GWAS), demonstrating the potential importance of splicing misregulation in complex traits [22]. Previous work from our lab and others have shown that gene-by-environment interactions can impact both gene expression and complex traits [23, 24, 25, 26, 27, 28]. While splicing QTLs have been identified both in humans and mice [22, 29, 30, 31], less is known about how gene-by-environment interactions may affect RNA processing. The first step to address this question is to characterize RNA processing in response to environmental perturbations.

RNA processing is regulated in response to certain environmental stimuli, such as cancer therapy drugs, nutrient starvation and infection [32, 33, 34, 35] some of which influence cell viability [36, 37, 38]. For example, UV exposure leads to differential isoform usage in the gene *BCL2L1*, which is involved in the regulation of apoptosis. UV leads to increased abundance of Bcl-x_*s*_ which favors apoptosis as opposed to Bcl-x_*l*_ which is anti-apoptotic [39]. Other studies have demonstrated widespread, directed changes in the regulation of RNA processing. Infection with *Listeria monocytogenes* and *Salmonella typhimurium* led to increased inclusion of cassette exons and shorter 3’UTRs genome-wide [35]. The longer versions of 3’UTRs that were shortened were found to be enriched with particular microRNA binding sites, suggesting that the RNA processing shift leading to shorter 3’UTRs may be a way for these genes to evade down-regulation following infection. Despite the fact that these studies have increased our understanding of factors that influence changes in RNA processing, they have investigated only a limited number of environments. Cataloguing and characterizing RNA processing changes across many environments, in a tightly controlled study using specific treatments, is necessary to increase our understanding of the cellular mechanisms leading to variation in RNA-processing, including which aspects are common across many environments and which are specific to certain perturbations.

Our study aimed to systematically assess the impact of a broad range of environmental perturbations on the regulation of RNA processing. We measured RNA processing patterns in five cell types across over 30 treatments, corresponding to a total of 89 cellular environments with at least 3 biological replicates and additional technical replicates (297 RNA-seq libraries in total with 130M reads on average) [23]. The treatments represent compounds to which we are exposed in daily life, ranging from metal ions and vitamins to allergy medication. This work demonstrates the extent of alternative RNA processing in response to a wide range of specific environmental perturbations, indicating molecular mechanisms by which trans factors influence this process.

## Results

### External stimuli induce environment-specific shifts in RNA processing

Using high-throughput RNA sequencing, we identified 32 compounds that induce gene expression changes in 32,451 genes in 5 different cell types (a total of 89 environments) [23] (Table S1). In order to identify changes in RNA processing, we utilized the probabilistic framework implemented in the software Mixture of Isoforms (MISO) [40], which characterizes changes in exon usage by calculating a percent spliced in (PSI, Ψ) value. The Ψ value is calculated by quantifying the fraction of reads specific to an inclusion isoform, specifically, reads aligning to the alternative exon or its junctions. Instead of entire isoforms, which may involve multiple RNA processing mechanisms that are convolved together, we focused on exons that are tied to known RNA processing mechanisms. We focused on events that involve known curated isoforms (see methods), rather than novel isoforms, and characterized variation in RNA processing events across different environments. This allowed us to learn about cis- and trans-acting mechanisms leading to the RNA processing response. Specifically, we characterized changes in eight event types: skipped exons (SE), retained introns (RI), alternative 3’ or 5’ splice sites (A3SS, A5SS), mutually exclusive exons (MXE), alternative first or last exons (AFE, ALE), and tandem untranslated regions (TandemUTR) (Figure 1A, Supplementary File 1 - Figure S1 shows a treatment color key used throughout the manuscript). Across all conditions, we identified 15,300 changes in RNA processing, representing a unique set of 8,489 events that significantly differ between at least one treatment and control conditions (Table 1, Supplementary File 2). These events are found in genes enriched for gene ontology terms such as RNA binding, gene expression, metabolic process, response to stress and cell cycle suggesting their role throughout the cellular response to environmental perturbation (BH *p*-value *<* 0.05, Supplementary File 1 - Table S2) [41]. Each significant change in RNA processing event was identified based on RNA sequencing data across cell lines derived from three unrelated individuals (example in Figure 2 and at http://genome.grid.wayne.edu/RNAprocessing). Across all environments, the most abundant event types with shifts were RI, AFE and ALE (relative to the number of sites tested), while the least abundant was A3SS (Figure 1B).

**Figure 1:**
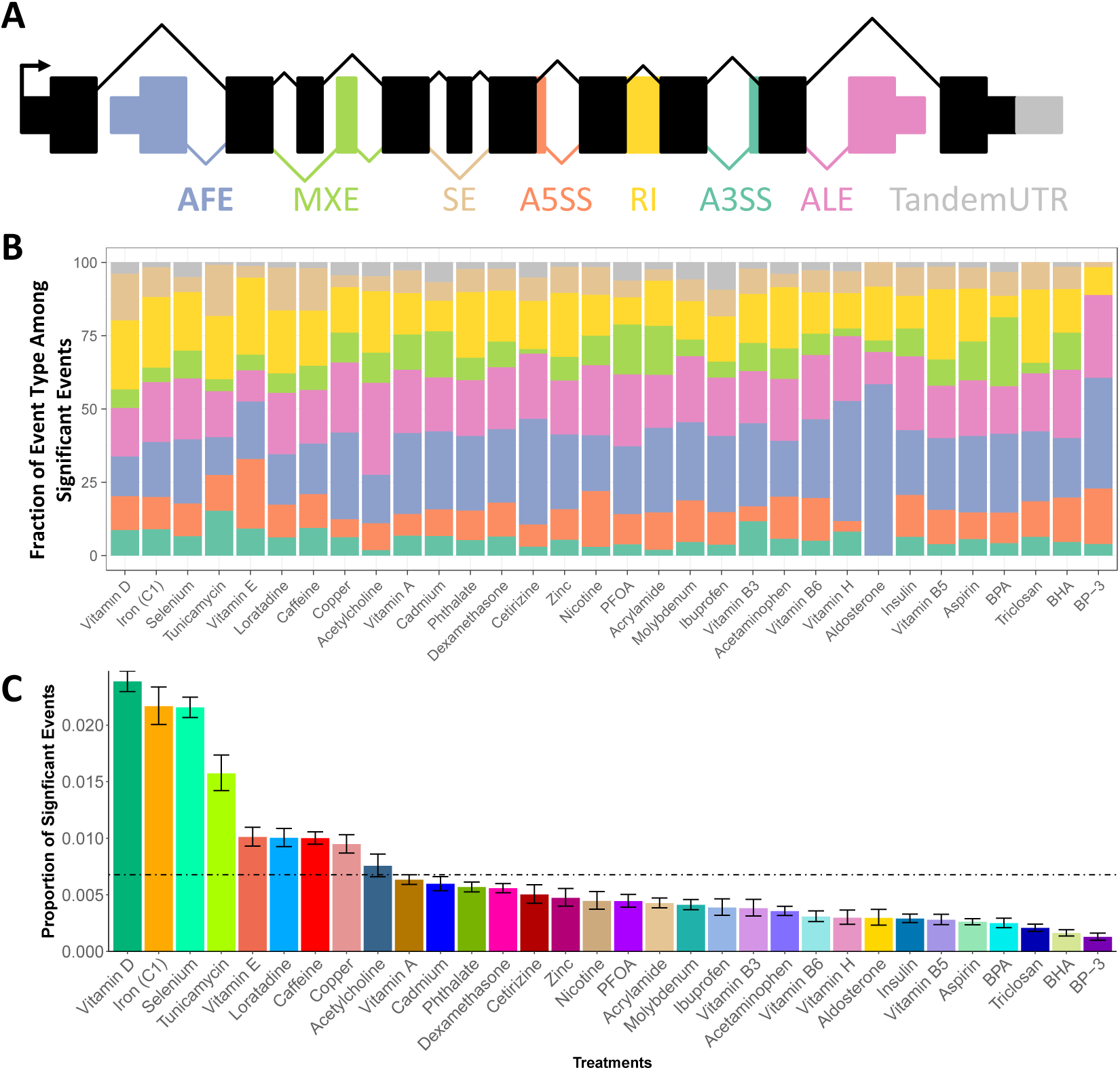
RNA Processing Events and Gene Expression Changes Following Treatment. A) Diagram depicting the 8 types of RNA splicing changes characterized in this work: alternative first exon, mutually exclusive exon, skipped exon, alternative 5’ splice site, retained intron, alternative 3’ splice site, alternative last exon, and tandem untranslated region. B) Graph showing the estimated proportion of each event type within a given treatment resulting from a logistic model. C) Proportion of significant changes in events over the total number of events that were tested in that treatment. Each bar combines all cell types treated with the compound. Error bars denote the standard error from a binomial test. The dotted line indicates the average proportion of significant events across all treatments.

**Figure 2:**
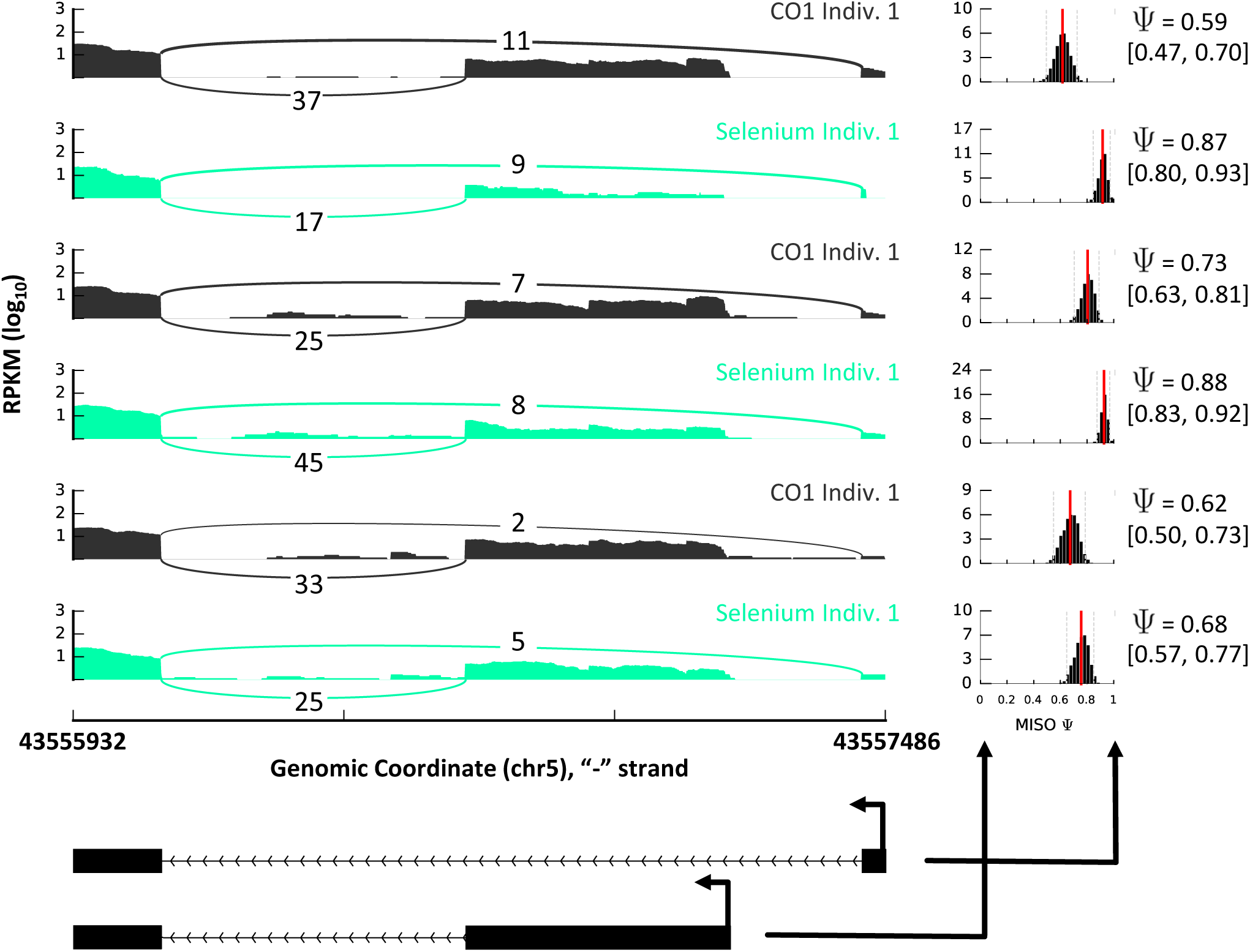
Sashimi plot showing AFE change following selenium treatment in LCLs across 3 unrelated individuals. The plots on the right show the Ψ value for each sample with confidence intervals. The plots to the left show the read coverage in each exon with model of this region below. High Ψ indicates preference for the upstream AFE.

**Table 1:** RNA processing events across 89 environments.

|  | SE | A3SS | A5SS | RI | MXE | AFE | ALE | TandemUTR |
| --- | --- | --- | --- | --- | --- | --- | --- | --- |
| # of sig. changes | 1681 | 516 | 686 | 1759 | 669 | 6015 | 3726 | 248 |
| # of sig. events | 1144 | 354 | 370 | 960 | 404 | 3003 | 2075 | 179 |
| # of events tested | 15895 | 5357 | 3714 | 3731 | 3728 | 10926 | 6714 | 2335 |
| % sig. events | 7.2 | 6.6 | 10.0 | 25.7 | 10.8 | 27.5 | 30.9 | 7.7 |

As we studied events in a given cell type across conditions, we found treatment-specific shifts in RNA processing, resulting in vast differences in the number and type of event shifts (examples in Figure 3). We found a wide range in the number of significant shifts across treatments, with vitamin D producing the highest number (2,530 events) and BP3 leading to the lowest number of significant shifts in RNA processing (65 events) (average = 478 events, 0.5% of events tested) (Figure 1C). The number of RNA processing changes in each environment is minimally correlated to the sequencing depth of the library (Spearman’s *ρ* = 0.15, *p* = 0.02; Figure S2A), but it is correlated to the number of differentially expressed genes in each environment (Spearman’s *ρ* = 0.59, *p* = 1.22 × 10^−8^; Supplementary File 1 - Figure S2B), suggesting the same underlying mechanism inducing changes in RNA processing and in overall gene expression.

**Figure 3:**
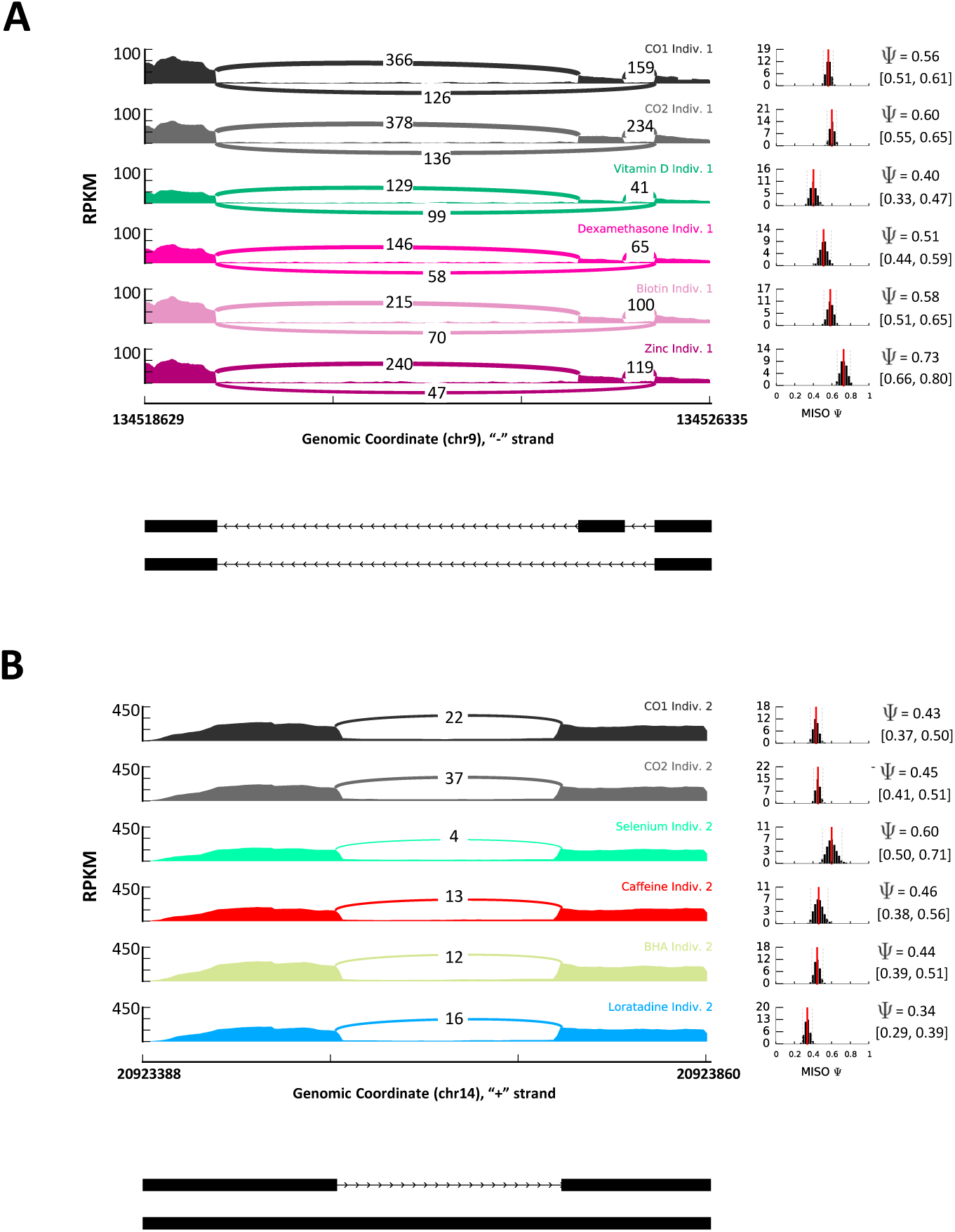
Example of RNA Processing Across Treatments in One Individual. The plots on the right show the Ψ value for each sample with confidence intervals. The plots to the left show the read coverage in each exon with a model of this region below each read coverage plot. A) Sashimi plot showing SE shift in *RAPGEF2* in PBMCs following exposure to 4 treatments (vitamin D, dexamethasone, biotin and zinc) and both controls. B) Sashimi plot showing SE shift in *APEX1* in melanocytes following exposure to 4 treatments (selenium, caffeine, BHA and loratadine) and both controls.

In addition to differences in the overall number of splicing changes, we also found differences in relative number of changes in certain event types (Figure 1B). While changes in AFEs represent the greatest overall number of changes across environments, there is substantial variation in the extent to which each event type changes within each treatment (Figure 1B). We utilized a generalized linear model to determine the proportion of event types among significant event shifts in a given treatment. With this model, we identified 3 treatments that showed enrichment for an event type, including vitamin E, tunicamycin, and cadmium. For example, vitamin E is enriched for A5SS while cadmium is depleted for RI (Figure 1B). Together, these results demonstrate that, similar to changes in gene expression, a large number of RNA processing events change in response to environmental perturbations.

### Direction of RNA Processing Shifts and Gene Expression Changes

Regulation of RNA processing events in response to environmental perturbations may be mediated by trans factors that impact many RNA processing events of the same type, or by cis-acting regulatory sequences that would impact each event separately. To investigate these two mechanisms we considered global shifts in RNA processing. Specifically, among the 8 event types with changes following treatment, 5 can be thought of directionally: SE, RI, AFE, ALE, and TandemUTR. Each event can be given a sign such that a positive ΔΨ (same as positive Z-score) means an increase in usage of the skipped exon, upstream AFE, downstream ALE, longer TandemUTR or intron retention in the treatment sample as compared to control (Figure 4A).

**Figure 4:**
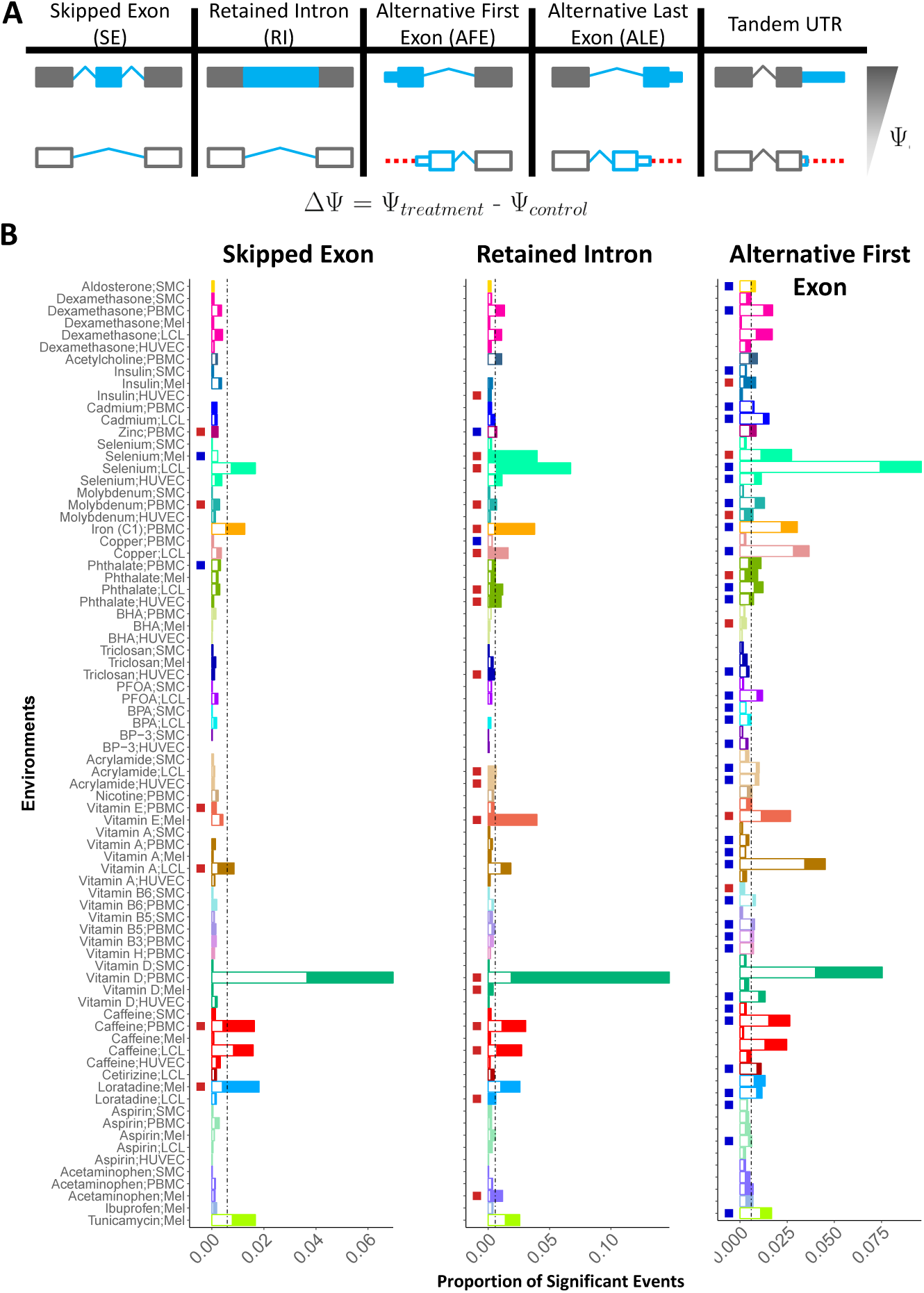
Direction of Shift in Events Following Treatment. A) Schematic of direction of event shifts for a given Ψ and ΔΨ. Shown for 5 event types. B)These plots indicate the direction of shift for 3 event types: SE (left), RI (middle) and AFE (right). Each plot shows 78 environments for which these events were tested. The height of each bar shows the proportion of significant event shifts for each environment. Each bar is then broken in two with the shaded region showing the proportion of the significant changes that shifted towards a positive ΔΨ (inclusion of exon, intron or upstream AFE) while the white region of each bar is the proportion of sites with a negative ΔΨ. The column of boxes shows if there is a departure from the expected 50:50 for positive to negative ΔΨ (tested using a binomial test). Red denotes enrichment for ΔΨ *>* 0 and blue for ΔΨ *< 0*).

When we focused on treatments with at least 30 significant RNA processing shifts of a certain event type, 19% of treatments showed a correlation (*p <* 0.05) between changes in RNA processing and changes in gene expression (examples in Supplementary File 1 - Figure S3, Table S3). For example, iron induced a positive correlation between ALE and gene expression (Spearman’s *ρ* = 0.27, *p* = 0.002) (Supplementary File 1 - Figure S3A). Specifically, genes shifting towards usage of the downstream ALE following iron treatment also have increased expression in the treatment samples. On the other hand, selenium leads to the opposite effect: increased expression following selenium is found in genes that utilize the upstream ALE (Spearman *ρ* = − 0.18, *p* = 1 × 10^−4^) (Supplementary File 1 - Figure S3B). These data suggest that cells respond to specific environmental perturbations with concerted shifts in RNA processing events and gene expression.

### Coordinated RNA Processing Shifts Across Cellular Environments

We investigated whether the global shifts in events had consistent direction across environments suggesting a shared trans-acting mechanism of change. First, we found that 8 environments led to an enrichment for SE shifts toward either inclusion or exclusion of the alternative exon (Figure 4B). Specifically, six environments were enriched for positive SE shifts which indicate global inclusion of the alternative exon while two led to more negative shifts or exclusion of the exon.

When studying RI across all environments, we identified 20 environments that lead to global shifts in intron inclusion. Specifically, 18 out of 20 were enriched for positive events (as compared to the expected proportion of 50%) (Figure 4B), thus showing enrichment for intron retention as compared to the control for most environments. These results suggest a common mechanism for intron retention in cells that respond to changes in the environment. For example, even though vitamin D causes many more changes in alternative splicing in PBMCs, all cell types trend towards retaining introns following vitamin D treatment. This can be more clearly seen when considering all RI events (not just significant events), where all 4 cell types show a shift toward more positive values in their ECDF (Kolmogorov-Smirnov (KS) test *p <* 0.05 for 3 of 4 cell types) denoting higher ΔΨ values, retaining of introns, following vitamin D treatment (Supplementary File 1 - Figure S4A and B).

Of the 5 event types whose direction could be assessed, AFE global shifts are observed in the most environments (Figure 4B). Specifically, 34 environments led to usage of the downstream AFE (negative Z-scores), while seven treatments were significantly enriched for shifts to the upstream AFE (positive Z-scores) (Figure 4B). Interestingly, several treatments lead to opposite AFE shifts in different cell types demonstrating the importance of the cellular background in response to environmental perturbations. For example, insulin leads to a shift toward the downstream AFE in SMCs but a shift toward the upstream AFE in melanocytes. This is also apparent when we consider all event shifts in these environments (KS test *p <* 0.05) (Supplementary File 1 - Figure S4C and D). Both ALE and TandemUTR also showed deviation from the expected 50:50 ratio of positive to negative events but the trend was less clear (Supplementary File 1 - Figure S5). These results demonstrate that global shifts in RNA processing events can be determined solely by the treatment or by the combined effect of treatment and cell type.

### Environmental shifts in SE and RI are mediated by changes in splicing factors expression and binding

In order to elucidate the specific factors involved in the global shifts in SE and RI events, we focused on factors likely to influence RNA processing, specifically splicing factors. We quantified the gene expression changes of splicing factors across all environments to determine if there was a correlation to the number of positive (inclusive) RNA processing shifts. The underlying hypothesis is that shifts in exon usage may be explained by splicing factors that: 1) have activity largely mediated by changes in gene expression, and 2) have the same influence over splicing in all treatments.

We identified 14 splicing factors (of 166 tested) with changes in gene expression correlated with percent positive events for RI (BH FDR *<* 5%, example in Figure 5A and B, Supplementary File 1 - Table S4); none were found for SE. Notably, we identified that changes in expression of *LARP7* are positively correlated with RI events. This suggests that the increased expression of *LARP7* under treatment conditions leads to more intron retention (positive RI events). Previous work has shown that *LARP7* promotes skipping of alternative exons [42]. Our results suggest that *LARP7* also plays a role in intron retention events. This trend can be seen across all environments considered. For example, selenium leads to an increase in expression of *LARP7* and more intron retention.

**Figure 5:**
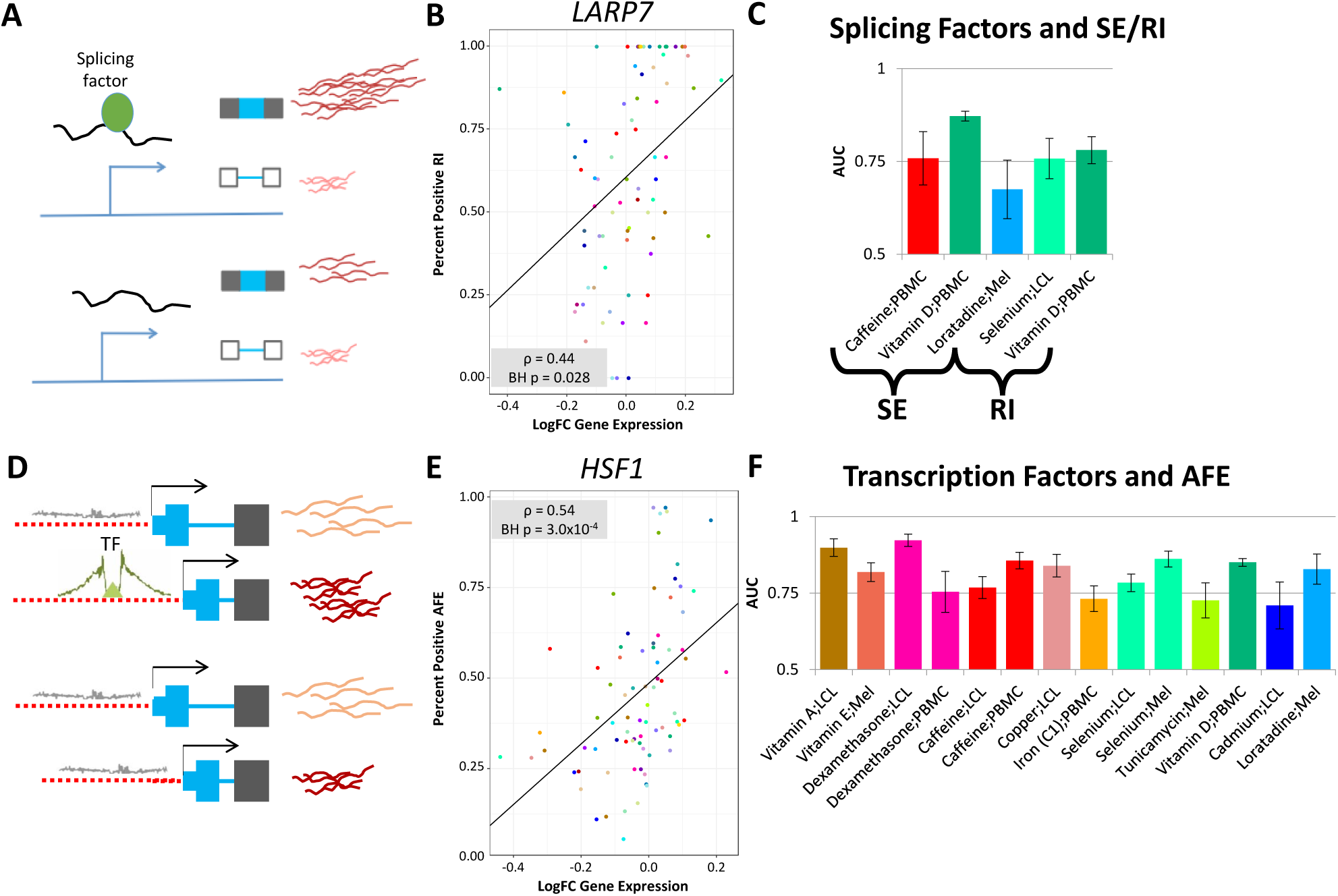
Effect of Trans-Factor Binding on RNA Processing Shifts. A) and D) show models of hypothesized mechanism of splicing or transcription factor influence on RNA processing and exon usage. B) An example of a correlation between the changes in gene expression of an RNA processing factor (*LARP7*) and the percent of RIs that shift towards the intron retention across all environments for which gene expression could be assessed. E) An example of a correlation between the changes in gene expression of a transcription factor (*HSF1*) and the percent of AFEs that shift towards the upstream AFE across all environments for which gene expression could be assessed. The correlation for B) and E) was tested using Spearman’s rho and the p-value shown is Benjamini-Hochberg corrected while the trendline depicts the best-fit line. C) Graph indicating the predictability (AUC as a proxy) of SE or RI shifts in a certain environment given predicted splicing factor binding sites (RNAcompete). F) Graph indicating the predictability (AUC as a proxy) of AFE shifts in a certain environment given transcription factor footprints [48].

Many factors may influence RNA processing differently following various treatments and so we may miss an effect by investigating common expression patterns across environments. Also, some factors are known to have different effects depending on binding location and not necessarily on overall gene expression. For example, when SR proteins (serine-arginine proteins) bind upstream of 5’ splice site, they induce splicing but do not have the same effect when bound in the intron [43]. With this in mind, we asked whether predicted binding sites of splicing factors may explain SE and RI. First, we characterized motifs that are present upstream, downstream or within the alternative unit (exon for SE or intron for RI). We, then, utilized an elastic-net regularized generalized linear model (GLM-NET) to predict splicing changes in 5 environments with greater than 100 significant event shifts (3 with SE, 2 with RI), based on the binding motif occurrences of splicing factors. When studying the model as a whole, we found that area under the curve (AUC) for each environment ranges from 0.67 for melanocytes exposed to loratadine to 0.87 for PBMCs exposed to vitamin D, suggesting that binding of splicing factors is important for determining changes in splicing following treatment, but the impact differs across cellular environments (Figure 5C). We also found that the genomic location of a binding site, relative to the splicing event, is an important predictive feature. For example, a motif for *RBM8A* (M054 0.6 from RNAcompete [44]) is a part of the predictive model of SE in PBMCs treated with vitamin D but only when the motif is located in the upstream intron. This demonstrates the positional effect of binding that others have characterized for some splicing factors [43, 45, 46, 47] and expands its importance across a large number of environmental perturbations.

### Effect of transcription factor expression and binding on AFE

We hypothesized that transcription factors regulate AFE shifts and TSS usage in response to environmental perturbations. Similar to our analysis with splicing factors, we first hypothesized that shifts in AFE could be the consequence of changes in gene expression for transcription factors that promote usage of either the upstream or the downstream TSS and have similar effects in all environments.

We identified 328 (out of 1,342) transcription factors whose change in expression is correlated with shifts in AFE (BH FDR *<* 5%) (example in Figure 5D and E, Supplementary File 1 - Table S5). Together, these results suggest that transcription factor binding influences the choice of TSS leading to a consequent shift in alternative first exon usage.

To directly determine the effect of TF binding on AFE shifts, we then utilized transcription factor footprints identified in DNase-seq data from ENCODE and the RoadMap Epigenomics [49, 48, 50] to predict shifts in AFE usage in 14 environments. We used footprints from more than 150 cell types to better capture a wider range of cellular environments, as determined by tissue of origin or culturing conditions. To predict AFE shifts, we considered the number of footprints present within 1000bp in either direction of each transcription start site (defined as the beginning of each alternative first exon), and used GLM-NET (as we did in the splicing factor analysis). Across the 14 environments, the AUC ranges from 0.71 for cadmium in LCLs to 0.92 for dexamethasone in LCLs (Figure 5F). These data suggest that transcription factor binding predicts changes in AFE following treatment.

### Validation of the mechanism for AFE shifts

By inducing changes in transcription factor binding, specifically by perturbing the cellular environment, we can validate the effect of binding on AFE usage (Figure 6A). To this end, we performed ATAC-seq in LCLs following treatment with selenium and its vehicle control. First, we noticed that selenium leads to an overall reduction in chromatin accessibility near transcription start sites (Figure 6B). To determine how selenium influences binding of transcription factors near alternative TSS, we characterized chromatin accessibility following treatment with selenium, or control (with the footprints used in the prediction analysis).

**Figure 6:**
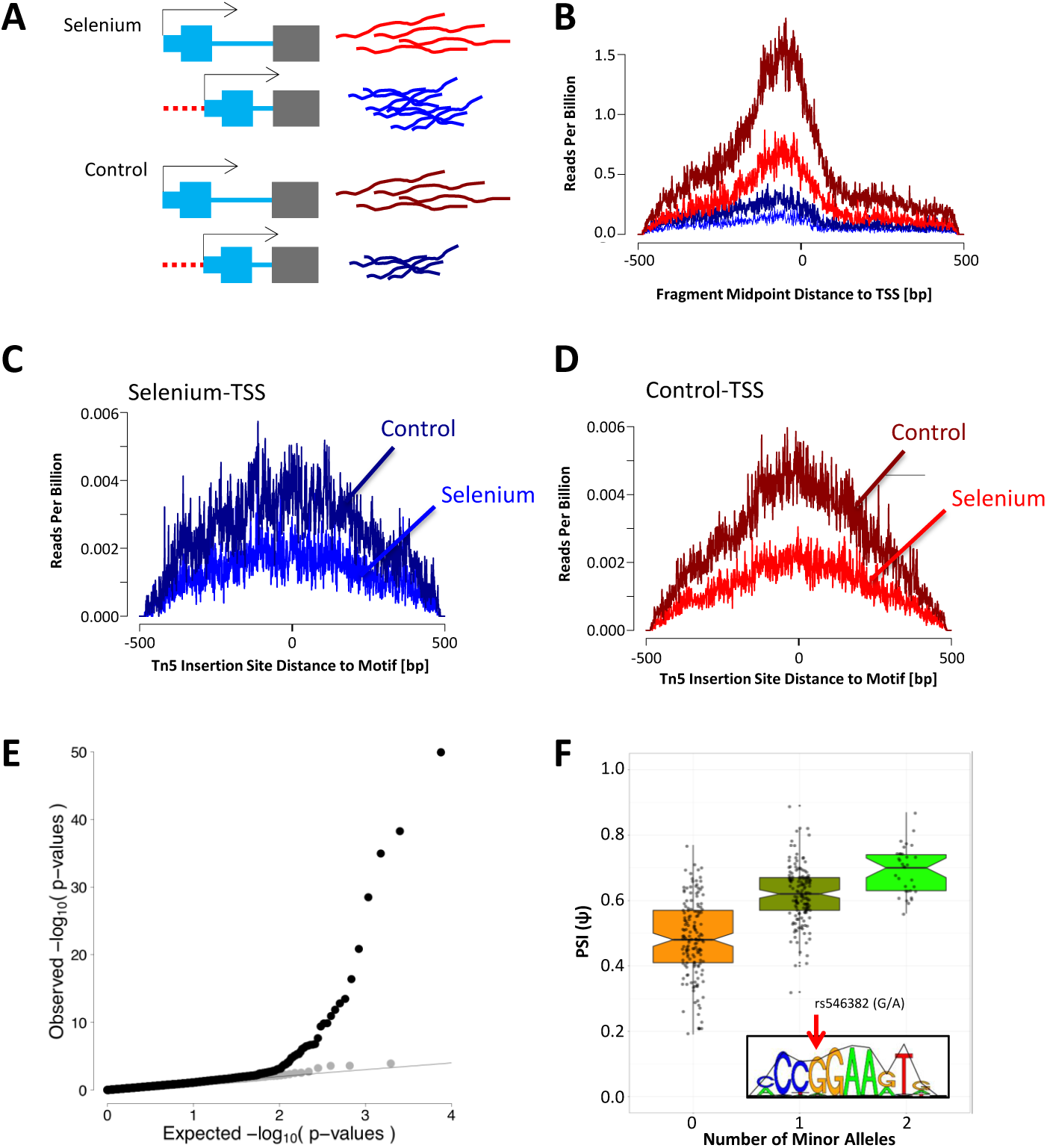
Binding of ELF2 Impacts AFE Shift Following Selenium Treatment. A) Model illustrating the situation where transcription is shifted towards the downstream AFE following selenium treatment and demonstrates an example of a TSS that would be included in our analysis. B) ATAC-seq denotes chromatin accessibility profile (read count normalized to total reads in the library) centered on TSS with AFE. Specifically, blue lines show accessibility at TSS to which the AFE shifts towards in selenium while red show the other TSS (as illustrated in A). Accessibility following selenium treatment is denoted by the bright blue and bright red lines while the control sample accessibility is denoted by the dark blue and dark red lines. C and D) ATAC-seq profiles centered on ELF2 motif locations within 1000bp of either TSS (colors are the same as in A and B). The difference in the ratio of treatment vs control read counts between the preferred and not preferred AFE is significant (BH p-value = 2.0 × 10^−3^). E) QQ-plot of the AFE QTL p-values for SNPs in footprint of ELF2 (within cis region of an AFE event) that are predicted to influence binding (black) or not (gray). F) Association between genotype of SNP (rs546382) found in an ELF2 footprint and predicted to influence binding [48] and Ψ of AFE in *IGHMBP2* across European individuals from the GEUVADIS data (*p*-value = 1.1 × 10^−35^). In the bottom right of the graph is the motif logo for ELF_2_ and the arrow indicates the position of the SNP.

We found significant differences in chromatin accessibility for 64 motifs, near the TSS that was preferentially used in the treatment versus the TSS preferred in the control condition. Of these 64 motifs, 26 are ETS transcription factor family members (or from motifs with similar sequence preferences). The most significant motif was for the transcription factor ELF2 (also in the ETS family, *p*-value = 5.4 ×10^−6^). We found a global decrease in chromatin accessibility at the ELF2 motif but there was a milder decrease in accessibility at the preferred TSS following selenium (Figure 6C and D), compared to the non-preferred TSS. These data suggest that at baseline ELF2 promotes transcription at both TSSs. However, following selenium treatment, though there is an overall decrease in ELF2 from both TSSs, there is a greater decrease from one TSS and this leads to a shift towards less usage of that TSS following treatment. All 26 motifs predicted from the ETS family of transcription factors show a similar change in binding as ELF2. More broadly, these results support a mechanism for changes in TSS usage driven by changes in chromatin accessibility and potentially transcription factor binding in response to perturbations of the cellular environment.

To further validate the effect of ELF2 binding on AFE usage, we characterized AFE across 373 unrelated, European individuals from the GEUVADIS data [51]. We identified 8,263 AFE events that can be characterized in at least 200 individuals. Using these data we performed AFE quantitative trait loci (QTL) analysis, by focusing on the SNPs in ELF2 footprints in the cis region of an AFE. We found an enrichment for QTLs in SNPs that were also predicted [48] to impact binding of ELF2, compared to those that do not affect binding (Fisher test *p*-value *<* 0.05, OR = 3.14, Figure 6E and F). In this way, using genetic perturbation, we were able to validate the impact of transcription factor binding, and specifically binding of ELF2, on AFE usage.

## Discussion

We describe 15,300 event shifts following a wide range of environmental perturbations at 8,489 unique RNA processing event sites. We have provided a browsable web-resource cataloguing these RNA processing shifts. Researchers interested in a given gene, isoform, or treatment will be able to access our data to determine when RNA processing shifts occur and which other genes respond under similar environments. Mining of our results has the potential to inform on the mechanisms by which a cell responds to environmental perturbations and its genome-wide effect on RNA processing.

The majority of events could be characterized as AFE, ALE, RI or SE, suggesting that these are the most influenced by the environmental perturbations considered here. This is distinct from previous reports that TandemUTR events change most following infection [35] and suggests diverse mechanisms through which the cells respond to their environment.

Previous work has studied the role of splicing factors and transcription factors in RNA processing, in the absence of specific environmental perturbations. For example, others have shown that multiple splicing factors influence cassette exon usage, several of which fall into 2 protein families: hnRNPs and SRSFs. These 2 protein families often result in opposite splicing patterns [43]. These proteins may play a role in several of the RNA processing events that we study here, including SE, RI, A5SS, and A3SS. There are other studies that characterized proteins related to polyadenlyation site usage (which we study as TandemUTR), including E2F, CSTF2, CSTF64 [52, 53]. Furthermore, recent studies have suggested that binding of transcription factors may influence differential use of transcription start sites in mice [54]. While these studies demonstrate the role of trans factors in RNA processing, we aimed to determine their role in global RNA processing changes in response to environmental perturbation.

Across 89 cellular environments, we found that binding sites for specific trans factors predict the shifts in events following treatment, thus demonstrating the importance of these factors and their binding locations for cellular response. We often find that not all binding sites for a given motif are predictive, but rather only binding sites in a certain location relative to the exon of interest. Furthermore, while previous studies have demonstrated the impact of binding location on RNA processing events at baseline, we demonstrate that the effect of binding in a certain location is treatment-specific. These results highlight the importance of studying trans factor binding across various environments. Further analysis of these binding sites will aid in understanding the details of the molecular mechanisms regulating RNA processing response to each cellular environment. For example, motifs associated with weaker binding of a trans factor may allow for more rapid changes in RNA processing and a more rapid cellular response.

Previous reports have characterized differences in transcription factor expression and binding across cellular environments [49, 55]. Here, we show that variation in transcription factor binding following environmental perturbations may determine TSS usage in addition to their function of influencing total gene expression. Feng *et al.* demonstrated the influence of transcription factors on TSS usage in mice [54]. Here, we expand on this knowledge by showing a similar function in human cells, both in response to many environmental changes and across individuals. Using ATAC-seq data, we further pinpointed factors such as ELF2 whose binding is disrupted by the environment, leading to changes in TSS usage. The changes in TSS usage are also observed when binding is disrupted by genetic variation in the GEUVADIS data, as shown in the AFE QTL analysis. Transcription factors are often regulated by environmental changes and are then responsible for impacting expression of many genes to promote re-establishment of cellular homeostasis (reviewed in [56]). Therefore, we suggest that TSS usage may also play a substantial role in cellular response and homeostasis.

Alternative RNA processing is predicted to occur in over 95% of multi-exon genes in humans across various tissues [57]. Our comprehensive catalog of genome-wide RNA processing changes can be utilized in future studies that aim to understand the role of RNA processing under various conditions and diseases as many of the treatments we used represent compounds to which individuals are commonly exposed. Furthermore, because RNA processing is associated with complex trait variation [22, 17], individual differences in RNA processing, specifically in response to environmental changes, could shed light on variation in organismal phenotypes.

## Methods and Materials

### RNA-seq data source

We used deep-sequenced RNA-seq data (fastq files) from Moyerbrailean *et al.*, 2016 [23]. Briefly, five cell types (LCL, PBMC, HUVEC, melanocyte and smooth muscle cells) were treated with 50 compounds to which humans are regularly exposed. Each environment (cell type and treatment) was represented in cell lines from three, unrelated individuals. We focused on 89 environments, with at least 80 differentially expressed genes, (Supplementary File 1 - Figure S1) that were sequenced to an average of 130M reads/library (297 RNA-sequencing libraries). These 89 environments include treatments and three vehicle controls (Supplementary File 1 - Table S1).

### Alignment

In order to detect alternative splicing, we used Mixture of Isoforms (MISO) [40], which requires reads of the same length. Therefore, we selected reads with a length greater than or equal to 120bp. All reads were trimmed to 120bp. We also removed reads whose paired end was less than 120bp.

Reads were aligned to the hg19 human reference genome using STAR [58] (https://github.com/alexdobin/STAR/releases, version 

~~~
STAR 2.4.0h1
~~~

), and the Ensemble reference transcriptome (version 75) with the following options:

~~~
STAR --runThreadN 12 --genomeDir <genome>
     --readFilesIn <fastqs.gz> --readFilesCommand zcat
     --outFileNamePrefix <stem> --outSAMtype BAM Unsorted
     --genomeLoad LoadAndKeep
~~~

where 

~~~
<genome>
~~~

 represents the location of the genome and index files, 

~~~
<fastqs.gz>
~~~

 represents that sample’s fastq files, and 

~~~
<stem>
~~~

 represents the filename stem of that sample.

For each sample (individual cell line with a given treatment), we merged sequencing replicates (across lanes and runs on the same sequencer) using samtools (version 2.25.0). We further removed reads with a quality score of *<* 10 (equating to reads mapped to multiple locations).

### Running MISO to detect splicing events

In order to detect alternative splicing, we used MISO on samples aligned as above. We utilized the events annotated and listed on http://miso.readthedocs.io/en/fastmiso/index.html. Specifically, we searched our data for 8 types of events with 5 from version 2 (SE, RI, A5SS, A3SS, and MXE, http://miso.readthedocs.io/en/fastmiso/annotation.html) and 3 from version 1 (AFE, ALE and TandemUTR, [59]). Two versions were used because AFE, ALE and TandemUTR were not annotated in version 2. We then ran miso.py on each of our samples for each of the 8 event types.

~~~
miso.py --run indexed_events/ my_sample1.bam --output-dir my_output1/
~~~

~~~
    --read-len 120
~~~

Then, we used 

~~~
summarize miso.py
~~~

 to get the summary statistics for each event in each sample, including the percent spliced in value (PSI, Ψ).

~~~
 summarize_miso --summarize-samples my_output1/ summaries/
~~~

To identify differential splicing, we used 

~~~
compare_miso.py
~~~

 which compares each event between treatment and control samples in the same individual cell line and experimental batch (plate).

~~~
 compare_miso --compare-samples my_output1/ my_control1/ comparisons/
~~~

This script resulted in a ΔΨ, a Bayes factor and p-value for each comparison. We then focused on comparisons where both treatment and control contained 2 reads covering each isoform uniquely and a total of 10 reads unique to either isoform for SE, RI, A5SS, A3SS, MXE, AFE and ALE. TandemUTR can only have reads specific to one isoform as the other isoform is simply a shorter version and completely overlaps the first. Therefore, we focused on comparisons of TandemUTR where both treatment and control contained 5 reads specific to the longer isoform and 10 total reads that covered either isoform.

Additionally, in order to inform on a cut-off for significant differential splicing, we performed comparisons between 2 controls (CO2 vs. CO1). Similar to treatments versus controls, we compared treatments performed in the same individual cell line and on the same plate. This generates an empirical null distribution that can be used to calibrate the statistical significance of the results. To this end, we used the same read requirements and filters as described above.

### Detecting significant changes in splicing

Because we had samples from three individuals for each environment (cell type and treatment), we aimed to combine the differential splicing scores across individuals. In order to do this, first we constructed tables for each event type that have all comparisons for all events of that type and take the Bayes factor (BF) computed by MISO. Next, we used our comparison of CO2 to CO1 to estimate the empirical null distribution. This is equivalent to an empirical null distribution used in permutation-based approaches to correct *p*-values. The empirical *p*-values for each treatment versus control BF are calculated by the corresponding quantile in the empirical null distributions of BF in the CO1 versus CO2 comparisons. This was done separately for each event type, as they may have different underlying distributions under the null hypothesis of no changes, i.e. ΔΨ = 0. Then, we converted each empirical *p*-value to a Z-score while retaining the direction of the change from the ΔΨ (calculated by MISO):

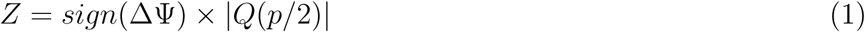

where *Q* represents the quantile normal function qnorm in R. These Z-scores are then added across all individuals with enough reads for the specific event considered in a given environment and divided by the square root of the number of individuals. We required that for a given isoform, at least 2 of the 3 individuals had high enough coverage to be measured (see read minimums in previous section). Finally, we ranked these new Z-scores (which are a combined measure across 2 or 3 individuals in an environment) and calculated a Benjamini-Hochberg (BH) corrected p-value to control for the false discovery rate (FDR). We considered an event with a significant shift if the BH FDR *<* 15%.

## Assigning direction to AFE and ALE

Of the 8 event types, 3 are tested directionally by MISO. SE, RI and TandemUTR had the isoforms assigned such that a higher Ψ corresponded to more inclusion of the skipped exon, inclusion of the retained intron or longer UTR, respectively. In order to assess directionality of ALE and AFE events, we modified the Ψ signs such that higher Ψ values corresponded to the more upstream AFE or downstream ALE (on the transcribed strand, using the transcription start site or end site, respectively) (Figure 4A). This was done for all analysis steps considering directionality.

## Gene expression changes

We calculated differential gene expression as described in Moyerbrailean *et al*. [23] using DESeq2 [60]. We analyzed the correlation between the number of significant shifts in exon usage (of any type) to genes that are differentially expressed in each environment using Spearman’s *ρ* (Supplementary File 1 - Figure S2).

In order to make a reasonable comparison to changes in gene expression, which are relative to the relevant control sample from the same cell type, we used the ΔΨ, which was calculated compared to the same control samples. For each treatment (including all relevant cell types), we found genes that had a significant shift in an RNA processing event and determined the correlation between log-fold gene expression changes (measured over all 3 cell lines with DESeq2) and the average ΔΨ across the same 3 individuals (Supplementary File 1 - Figure S3, and S3). In this analysis, we focused on treatment and event type combinations that had at least 30 RNA processing shifts in genes whose expression could be assessed in the same treatment across all cell types.

When we studied the correlation of a specific event to gene expression of either a splicing factor or transcription factor, we correlated the log-fold gene expression of that factor to the percent positive events (PPE) of a given event type across all environments:

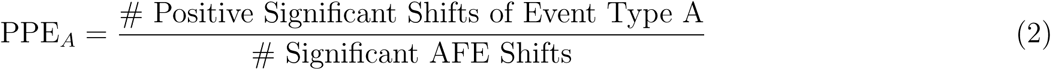

The correlation between PPE and log-fold gene expression is calculated using Spearman’s *ρ* and a best-fit line is added to each plot in Figures 5B and E.

### Enrichment of event types among significant events in each treatment

In order to estimate the fraction for each event type while controlling for cell-type, and to identify enrichment of a specific event type among events with significant shifts in a given treatment, we used a generalized linear model:

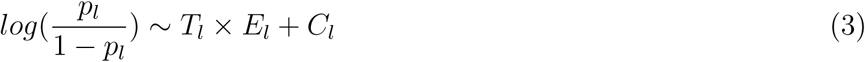

where *p*_*l*_ is the probability that the event ‘*l*’ has a significant shift of a specific type *E*_*l*_, for a given treatment *T*_*l*_ and cell-type *C*_*l*_. This allowed us to study the interaction between treatment and event type (while removing the effect of cell type) on whether or not an event type is significantly enriched for a specific treatment.

The model incorporates an intercept which utilizes the information from one of each category in the equation such that every *β*_*t,e*_ is relative to this baseline category. In order to create Figure 1B, we used the estimated fractions *p*_*l*_ from this logistic model, and conditional on each treatment *t*. We report the probability of each event type *e* among significant events. This allows us to compare proportion of event types across treatments. In order to determine enrichment of an event type among significant event shifts for a given treatment, we look at the *p*-value for the interaction term of treatment by event type (*β*_*t,e*_ ≠ 0).

### Gene ontology analysis

We utilized GeneTrail [41] to find enrichment of gene ontology terms. We compiled a list of unique genes that had significant changes in RNA processing. We focused on the 3,363 genes to which an RNA processing event could be uniquely assigned. We then determined which GO categories were over-represented as compared to a list of all genes to which an RNA processing event could be assigned. We considered a category over/under-represented if the Benjamini-Hochberg FDR *<* 5%.

### Elastic-net regularized generalized linear model

In order to assess the predictive power of transcription factor binding on RNA processing changes following treatment, we utilized the ‘glmnet’ package in R [61]. This package uses an elastic-net regularized generalized linear model (using a logistic link function). We used this model to assess the role of transcription factor binding on AFE and splicing factor binding sites on SE and RI in environments with at least 100 significant event shifts (BH FDR *<* 15%).

For our analysis of AFE, we used the transcription factor footprints derived from data collected by ENCODE and RoadMap Epigenomics [49, 48, 50]. We utilized footprints from all cell types because we expect binding to change following treatment and so did not want to be too restrictive on what we called a binding site. We then counted the number of footprints within 1000bp (upstream and downstream) of each TSS for each AFE that showed a significant shift following treatment. Next, for each AFE, we subtracted the number of footprints near the downstream TSS from the number of footprints near the upstream TSS. We then used these values, for each motif, as predictors in the model. The variable we attempted to predict using this model was the direction of the shift (ΔΨ) following treatment. From the results of glmnet, we then used the lambda.1se, the lambda that was 1 standard error from the lambda that resulted in the highest AUC, to define an AUC for AFE in a given treatment. Measurement of AUC and selection of lambda was done using the cross-validation procedure implemented in glmnet (

~~~
cv.glmnet
~~~

).

For our analysis of SE and RI, we used the splicing factor binding sites predicted from RNAcompete [44]. We split each SE or RI with a significant shift following treatment into 5 regions around the event site. For SE, we had the region of the upstream intron, the skipped exon itself, the downstream intron, 100bp upstream of the 3’ splice site, and 100bp downstream of the 3’ splice site. For RI, we used the region of the upstream exon, the intron in question, the downstream exon, 100bp upstream of the 5’ splice site, and 100bp downstream of the 5’ splice site. We determined whether or not a splicing factor motif was found in any of these regions in the same direction of transcription and then separately considered these into the model as predictors. The variable we attempted to predict using this model was the direction of the shift (Δ Ψ) following treatment. From the results of glmnet, we then used the lambda.1se, the lambda that was one standard error from the lambda that resulted in the highest AUC, to define an AUC for SE or RI in a given treatment. Measurement of AUC and selection of lambda was done using the cross-validation procedure implemented in glmnet (cv.glmnet).

### ATAC-seq in LCLs exposed to selenium

The lymphoblastoid cell line (LCL) GM18508 was purchased from Coriell Cell Repository. LCLs were cultured in serum containing charcoal-stripped FBS and treated for 6 hours with 1μM selenium as described in [48]. Cells were also cultured in parallel with the vehicle control (water), to represent the solvent used to prepare the treatment. We then followed the protocol by [62] to lyse 25,000-100,000 cells and prepare ATAC-seq libraries, with the exception that we used the Illumina Nextera Index Kit (Cat #15055290) in the PCR enrichment step. Individual library fragment distributions were assessed on the Agilent Bioanalyzer and pooling proportions were determined using the qPCR Kapa library quantification kit (KAPA Biosystems). Library pools were run on the Illumina NextSeq 500 Desktop sequencer in the Luca/Pique-Regi laboratory. Barcoded libraries from three replicates (25,000, 50,000 and 75,000 cells each) were pooled and sequenced on multiple sequencing runs for 100M 38bp PE reads.

Reads were aligned to the reference human genome hg19 using bwa mem ([63] http://bio-bwa.sourceforge.net). Reads with quality *<*10 and without proper pairs were removed using samtools (http://github.com/samtools/).

To assess global shifts in accessibility, reads with different fragment length were partitioned into four bins: 1) [39-99], 2) [100-139], 3) [140-179], 4) [180-250]. For each fragment, the two Tn5 insertion sites were calculated as the position 4bp after the 5’-end in the 5’ to 3’ direction. Then for each candidate motif, a matrix ***X*** was constructed to count Tn5 insertion events: each row represented a sequence match to motif in the genome (motif instance), and each column a specific cleavage site at a relative bp and orientation with respect to the motif instance. We built a matrix 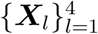 for each fragment length bin, each using a window half-size S=150bp resulting in (2×*S* + *W*) × 2 columns, where W is the length of the motif in bp. The motif instances were scanned in the human reference genome hg19 using position weight (PWM) models from TRANSFAC and JASPAR as previously described [64]. Then we used CENTIPEDE and motif instances with posterior probabilities higher than 0.99 to denote locations where the transcription factors are bound.

### Validating AFE mechanism with ATAC-Seq

First, we assessed chromatin accessibility within 500bp (in either direction) of each AFE by quantifying the number of reads (lengths 30-140bp, as this corresponds to lengths shorter than those that wrap around a nucleosome allowing us to focus on open chromatin). In order to obtain a sufficient number of sites, we collected all event shifts with BH FDR *<* 25%.The read count was summed across AFEs that were either upstream or downstream and then normalized to the total reads in the library (either selenium- or control-treated).

In order to compare the effect of transcription factor binding changes on AFE shifts following LCL exposure to selenium, we started with the transcription factor footprints from [48] used for the prediction analysis. We split these footprints into those found within 500bp of the TSS towards which transcription shifted following selenium versus those found within 500bp of the TSS which was less preferred following treatment (termed Selenium-TSS or Control-TSS). For this analysis, we studied all AFE shifts with a BH FDR *<* 25%. We used a more relaxed threshold for this analysis because we also must require the AFE to be within 500bp of a transcription factor footprint and would otherwise not have a sufficient number of sites to draw any conclusions. We then quantified the read counts in the selenium and control treated samples within 100bp (in either direction) of each of these motif locations and normalized these counts to the overall read counts in each library. For each footprint, we calculated the ratio of normalized read counts in treatment versus control libraries. We then used these ratios to perform a Student’s *t*-Test across all footprints of a specific transcription factor near the Selenium-TSSs compared to the Control-TSSs (2-tailed). Changes in transcription factor binding activity were considered significant if BH FDR *<* 5%.

### Validating effect of transcription factor binding with QTL analysis

We downloaded the bam files for 373 European individuals sequenced in GEUVADIS dataset [51]. We subsetted the sequencing data to obtain full-length reads of at least 75bp. Then, using the same parameters described above, we ran MISO to characterize AFE in these individuals. We quantified the *Psi* value for 13,712 unique AFE events (average of 8,495 per individual, max: 10,679 events, min: 5,567 events). We then normalized these *Psi* values across individuals by removing the effect of the lab that performed the RNA-sequencing, the population effect, and the first five principal components. Finally, we quantile normalized the resulting values, across the individuals that could be assessed. We focused on the AFEs that could be assess in at least 200 of the 373 individuals, leaving 8,263 events (also removing events on the X or Y chromosome).

To assess the impact of ELF2 binding on AFE usage, we used the annotation from Moyerbrailean *et al.* [65]. We focused on AFE events with SNPs in ELF2 footprint within 10Kb of each TSS(the first base of the event’s first exons).The association between genotype and normalized phenotype was measured using a standard linear model. For each SNP we assessed whether or not they were predicted to influence ELF2 binding [65]. We then compared the enrichment for QTLs among SNPs that impact ELF2 binding, as compared to those that are not predicted to impact ELF2 binding based on the sequence model derived from CENTIPEDE (i.e. the position weight matrix, PWM).

## Additional Files

### Supplementary File 1 — Supplemental Results

Supplementary text for additional results, figures and tables.

### Supplementary File 2 — 15,300 RNA Processing Shifts

This table describes all 15,300 RNA processing shifts that we identify in our data. These sites can also be found on our browsable web-resource (http://genome.grid.wayne.edu/RNAprocessing). This table has 20 columns as follows: 1) Unique identifier for each event, 2) Plate name which is a key covariate among our samples, 3) Event name from MISO database, 4) Chromosome of event, 5) Strand of mRNA, 6) Start positions for exons, 7) End positions for exons, 8) Treatment ID, 9) Treatment name, 10) Cell Type, 11) Control ID for the tested treatment, 12) Control name, 13) Type of event, 14) Number of individuals that could be assessed, 15) Number of individuals that had p-value derived from the logBF *<* 0.05, 16) Number of individuals that had positive ΔΨ and had p-value derived from logBF *<* 0.05, 17) Combined Z-score, q-value, 19) Ensemble gene ID, and 20) Gene symbol.

## Acknowledgments

We thank members of the Luca and Pique-Regi groups for helpful discussions and comments.

## Funding

This work was supported by the National Institutes of Health [5R01GM109215 to F.L and R.P.]; and the American Heart Association [14SDG20450118 to F.L.].

## Competing Interests

The authors declare that they have no competing interests.

